# Chaperonin-mediated Winter Cold Response via Circadian Clock Components in *Arabidopsis*

**DOI:** 10.1101/2024.05.09.593290

**Authors:** Goowon Jeong, Myeongjune Jeon, Jinseul Kyung, Daesong Jeong, Jinwoo Shin, Yourae Shin, Daehee Hwang, Youbong Hyun, Ilha Lee

## Abstract

Vernalization and cold acclimation are plant strategies that evolved to enhance their fitness during the winter season. The molecular mechanisms behind these processes in Arabidopsis have been intensively studied. However, how plants measure the duration of long-term cold exposure has not been fully understood. Our research shows that cytosolic chaperonin is crucial for appropriate long-term cold responses by regulating plant circadian clocks. Furthermore, one of the clock components, REVEILLE4 (RVE4) and RVE8 directly activate *VERNALIZATION INSENSITIVE 3* (*VIN3*) and the *C-REPEAT BINDING FACTOR* (*CBF*)/*DEHYDRATION-RESPONSIVE ELEMENT BINDING 1* (*DREB1*) family genes, which are key regulators of vernalization and cold acclimation, respectively. The activation of the *VIN3* and *CBF/DREB1* genes was specific to the time of day, indicating that cold exposure during the day is critical for long-term cold responses. Our research delves deeper into understanding the regulatory mechanism governing these two distinct long-term cold responses.

## Introduction

Plants, being sessile organisms, confront substantial survival and reproduction during the winter seasons, necessitating the evolution of two distinct strategies: vernalization and cold acclimation. Vernalization is a physiological process that enables biennial or winter annual plants to achieve flowering competence after winter cold. In *Arabidopsis*, the vernalization response is closely associated with the levels of a MADS-box transcription factor, FLOWERING LOCUS C (FLC), which suppresses flowering pathway integrator genes, *FLOWERING LOCUS T* (*FT*) and *SUPPRESSOR OF OVEREXPRESSION OF CONSTANS1* (*SOC1*), thereby preventing flowering in the fall. However, *FLC* levels gradually decrease during the winter season through epigenetic mechanisms (Sung and Amasino 2004). As a result, the pathway integrator genes are derepressed, and plants acquire the competence to flower in the upcoming spring (Moon et al. 2003). It is noteworthy that the *FLC* suppression is achieved over several weeks of cold exposure and is maintained during the next spring (Michaels and Amasino 1999; Sheldon et al. 2000). *VERNALIZATION INSENSITIVE 3* (*VIN3*), encoding a protein with a plant homeodomain (PHD), plays a key role in this epigenetic suppression. Its expression gradually increases during vernalization and the VIN3 protein recruits polycomb repressive complex2 (PRC2), leading to the accumulation of trimethylation of histone 3 lysine 27 (H3K27me3), inactive chromatin marks, on the nucleation region of *FLC* chromatin. In the next spring, H3K27me3 marks are spread over the entire *FLC* chromatin by LIKE-HETEROCHOMATIN PROTEIN1 (LHP1), leading to stable epigenetic suppression (Mylne et al. 2006). Currently, how long-term winter cold regulates *VIN3* expression is not fully understood, although recent studies have shown that transcription factors such as NAC WITH TRANSMEMBRANE MOTIF 1-LIKE 8 (NTL8), CIRCADIAN CLOCK-ASSOCIATED 1 (CCA1), LATE-ELONGATED HYPOCOTYL (LHY), and HEAT SHOCK TRANSCRIPTION FACTOR B2b (HsfB2b) are involved in its transcriptional regulation (Zhao et al. 2020; Jeong et al. 2022; Kyung et al. 2022). Null mutations in these genes, however, only cause minor changes in *VIN3* transcript levels after saturated vernalization.

Cold acclimation is a process by which plants increase their freezing tolerance after pre-exposure to low but non-freezing temperature (Thomashow 1999). It is caused by massive changes in the expression of cold responsive (*COR*) genes (Guy 1990; Fowler and Thomashow 2002). In *Arabidopsis*, three transcription factors, C-REPEAT BINDING FACTOR1 (CBF1)/DEHYDRATION-RESPONSIVE ELEMENT BINDING1B (DREB1B), CBF2/DREB1C, and CBF3/DREB1A, have been identified as upstream regulators of numerous *COR* genes, such as *RD29a/COR78* or *COR1512a* (Liu et al. 1998). Consistently, recent studies have shown that *cbf1 cbf2 cbf3* triple mutants show failure of cold acclimation (Jia et al. 2016; Zhao et al. 2016). Therefore, the three CBF/DREB1 proteins seem to be crucial for the freezing tolerance acquired through the cold acclimation. However, little is known about how CBF/DREB1 family proteins are regulated during long-term winter cold, thus enhances plant survival.

Meanwhile, previous studies have shown that the promoter regions of *VIN3* and *CBF/DREB1* genes (hereafter referred to as *CBFs*) contain evening element (EE) motifs, and two circadian clock transcription factors, CCA1 and LHY, play key roles in the expressions of *VIN3* and *CBFs (Fowler et al. 2005; Dong et al. 2011; Hepworth et al. 2018; Kidokoro et al. 2021; Kyung et al. 2022)*. The circadian clock in plants plays a crucial role in the appropriate response to diurnal changes in environmental signals, which are largely dependent on the presence and absence of sunlight. In *Arabidopsis,* the circadian clock consists of multiple feedback loops comprising many genes that cross-regulate each other, resulting in sustained daily oscillations in the expression of their target genes (Harmer et al. 2000; Fogelmark and Troein 2014). For example, CCA1 and LHY, two morning-phased MYB-related transcription factors, are known to repress many other clock genes such as *TIMING OF CAB EXPRESSION1*/*PSEUDO-RESPONSE REGULATOR1* (*TOC1/PRR1*), *PRR5*, *PRR7*, and *PRR9* as part of the feedback loops (Gendron et al. 2012; Nagel et al. 2015; Kamioka et al. 2016). On the other hand, unlike CCA1, which usually acts as a transcriptional repressor, other morning clock regulators such as REVEILLE (RVE) family proteins, at least five of 8 RVEs, RVE3, RVE4, RVE5, RVE6, RVE8, have been reported to act as transcriptional activators that directly bind to the EE motif on the promoter of evening-phased clock genes (Rawat et al. 2011; Hsu et al. 2013; Fogelmark and Troein 2014; Gray et al. 2017). Given that the effect of the *cca1 lhy* mutation on the vernalization response is minor, the molecular mechanism underlying the regulation by clock components during long-term cold exposure needs to be reevaluated (Kyung et al. 2022).

Chaperonins are protein complexes composed of several protein subunits, necessary for proper folding or stabilization of proteins (Kubota et al. 1995). They have a size of approximately 800-1,000 kDa and were previously classified into two paralog groups, Group I and Group II (Kubota et al. 1995; Leroux and Hartl 2000; Spiess et al. 2004). While Group I chaperonins were found in bacteria, chloroplasts and mitochondria, Group II chaperonins were found in archaea and cytosol of eukaryotes. TRiC (for <u>T</u>CP1 (Tailless-Complex Polypeptide 1) <u>Ri</u>ng <u>C</u>omplex), also known as CCT (for <u>C</u>haperonin <u>C</u>ontaining <u>T</u>CP1), is a Group II chaperonin that is required to stabilize cytoskeleton proteins and predicted to be necessary for the proper folding of many newly synthesized polypeptides, making it essential for cell survival in yeast and mammals (Horwich et al. 2007). TRiC consists of two sets of eight different subunits, called CCT1–CCT8, which form one ring structure, stacked back to back to create the whole TRiC complex (Cong et al. 2010). In *Arabidopsis*, TRiC is involved in the regulation of microtubules through its folding activity, and null mutations of CCT subunits have been shown to be lethal (Xu et al. 2011; Ahn et al. 2019), highlighting the essential role of CCT in plant cell viability. However, plant TRiC also has substrate specificity and plays a role in regulating developmental processes. For example, TRiC facilitates cell-to-cell trafficking of *KNOTTED1* to regulate stem cell function in the shoot apical meristem (Xu et al. 2011), suggesting that the role of TRiC is not limited to general housekeeping functions.

In this study, we have identified the upstream regulators of vernalization and cold acclimation during winter cold. We have discovered that the TRiC plays a crucial role in the optimal responses of vernalization and cold acclimation, which are mediated by circadian clock regulators, RVEs. We also showed that RVE4 and RVE8 directly bind to the EE motifs located in the promoters of *VIN3* and *CBFs*, revealing the genetic circuit involved in sensing winter cold. Our findings provide new insights into the signaling mechanisms that are mediated by the TRiC and circadian clock components during prolonged cold.

## Results

### A mutation in chaperonin causes reduced *VIN3* activation

To explore upstream regulators of *VIN3*, we screened 25,000 M2 mutant lines produced from the 3,412 M1 plants mutagenized with ethyl methanesulfonate (EMS) using a *VIN3p-GUS* reporter line (−0.2 kb *pVIN3_U_I::GUS*) (Kyung et al. 2022). From this screening, we isolated a mutant, *P161,* that shows a reduced GUS signal after 40 d of vernalization (40V) (Fig. 1A). This mutant also displayed growth retardation and pigment accumulation after 20V and 40V (Fig. 1B). The endogenous *VIN3* level in the mutant was also severely reduced compared to parental lines during vernalization treatment (Fig. 1C), confirming that *P161* carries a mutation in the gene required for *VIN3* activation.

**Figure 1.**
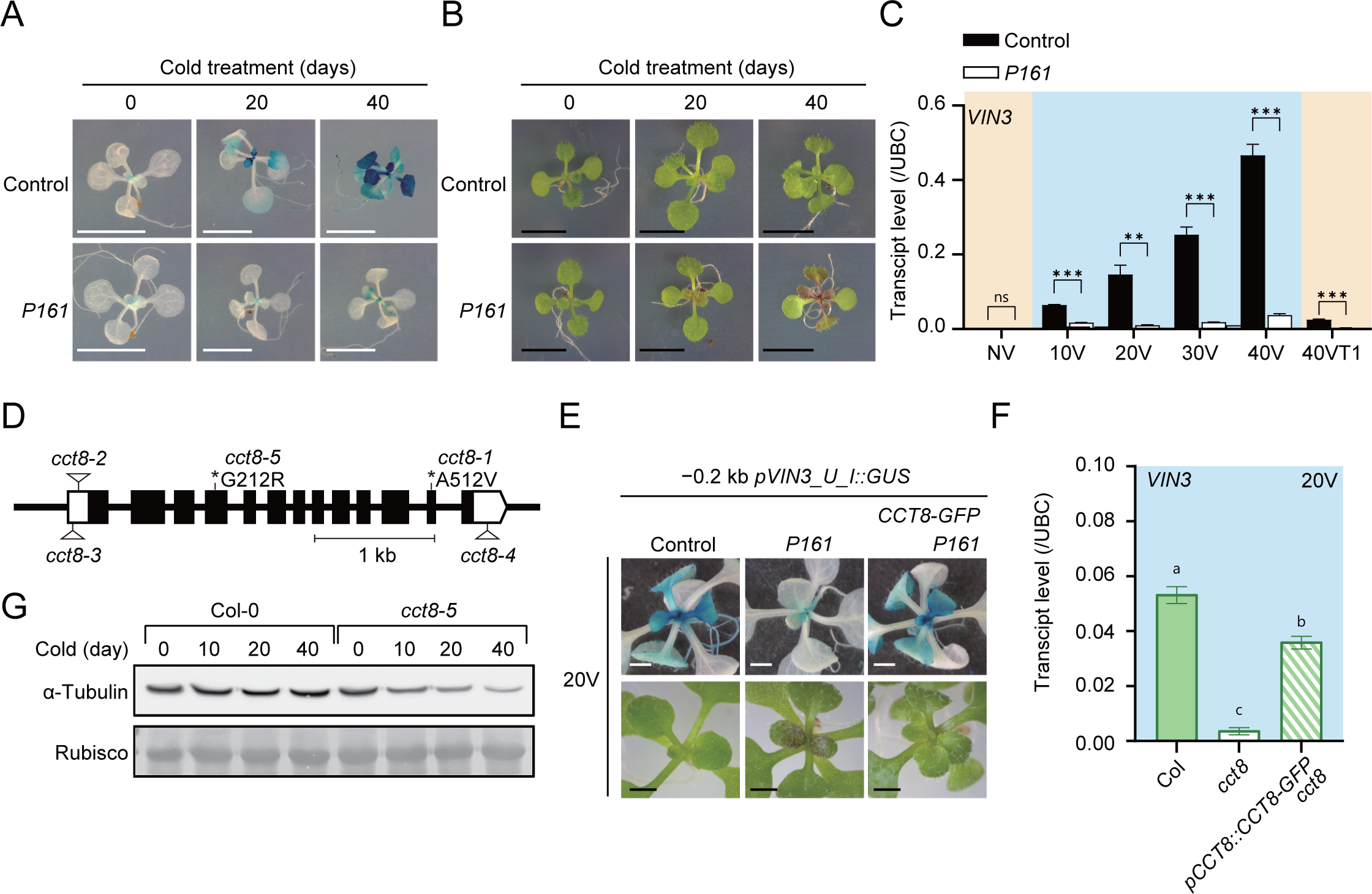
TRiC is involved in the *VIN3* activation. (A–B) Representative images showing the seedlings of parental lines (Control) and *P161*. The images show seedlings after (A) and before (B) GUS staining. Scale bars, 5 mm. (C) Comparison of endogenous *VIN3* transcript levels between parental lines and *P161* mutants during vernalization. Data are shown as means ± SEM for three biological replicates. Asterisks signify statistical significance (Student’s t-test; **p < 0.01, *** p <0.001, n.s., not significant). (D) Schematic illustration of the *CCT8* gene structure. Different mutant alleles are depicted by triangle or asterisks. Black boxes, white boxes, and black lines represent exons, untranslated regions, and introns, respectively. (E) Images of the seedlings of control, *P161*, and *P161* harboring *pCCT8::CCT8-GFP* transgene after 20V. The images show seedlings after (upper panel) and before (lower panel) GUS staining. Scale bars, 1 mm. (F) *VIN3* transcript levels in the seedlings of Col, *cct8-5* and *pCCT8::CCT8-GFP cct8-5* after 20V. Data are shown as means ± SEM for three technical replicates. Distinct letters indicate a significant difference as determined by one-way analysis of variance (ANOVA) with post-hoc Tukey test. (G) Immunoblot analysis revealing α-tubulin abundance in the seedlings of Col and *cct8-5* during vernalization. Rubisco was used as a loading control. NV, non-vernalized; 10V, 20V, 30V, and 40V indicate durations of vernalization for 10, 20, 30, and 40 days, respectively.

The F2 population, obtained from the cross between *P161* and parental line, showed a roughly 3:1 segregation ratio (290 WT vs 94 mutants, χ^2^=0.073), indicating a recessive mutation in a single gene locus. Through genome sequencing-based positional cloning (Lukowitz et al. 2000; Hou et al. 2010; James et al. 2013), we identified a missense mutation in At3g03960, coding for CCT8, a TRiC subunit, altering a conserved residue (Gly^212^ to Arg^212^) (Fig. 1D; Supplemental Fig. S1A,B). To confirm this, we checked *VIN3* levels in the previously reported *cct8-1* mutant (a substitution mutant, Ala^512^ to Val^512^) (Xu et al. 2011). Both *P161* and *cct8-1* showed similarly reduced *VIN3* levels and pigment accumulation after vernalization (Fig. 1E,F; Supplemental Fig. S2A), although *P161* showed relatively normal morphology compared to the *cct8-1* that exhibits upwardly curled-leaves and dwarfism. Consistently, the complementation test showed that *P161* is allelic with *cct8-1* (Supplemental Fig. S2B,C), and the introduction of the *pCCT8::CCT8-GFP* transgene into *P161* led to the recovery of vernalization-induced *VIN3* activation (Fig. 1E,1F). Taken together, our results indicate that the failure of *VIN3* induction in *P161* is caused by the missense mutation in the *CCT8.* Henceforth, we named the mutation *cct8-5*.

We checked if *cct8-5* also has a defect in chaperonin function for protein stability, because the *cct2* mutant has been reported to cause a defect in the chaperonin function for tubulin folding (Ahn et al. 2019). Although the *cct8-5* mutation did not significantly alter α-tubulin levels in NV, a gradual decrease in *cct8-5* mutants was observed over the vernalization period, contrasting with the minimal effect of vernalization on α-tubulin levels in the wild type (WT) (Fig. 1G). Therefore, it suggests that the lack of TRiC causes a cumulative effect on protein stability during vernalization.

### *cct8* mutation causes a defect in vernalization response

To analyze the effect of *cct8* on vernalization response, we introduced *cct8-5* into a vernalization-sensitive *FRI* Col line by genetic cross. The resulting *cct8-5 FRI* Col (hereafter *cct8 FRI*) flowered similarly to WT (*FRI* Col) without vernalization but showed delayed flowering after 10V and 20V (Fig. 2A,B). It suggests that the *cct8* mutation causes a defect in vernalization response at least in the early stages of vernalization. However, *cct8 FRI* exhibited a similar flowering time with WT after 30V and 40V (Fig. 2A,B).

**Figure 2.**
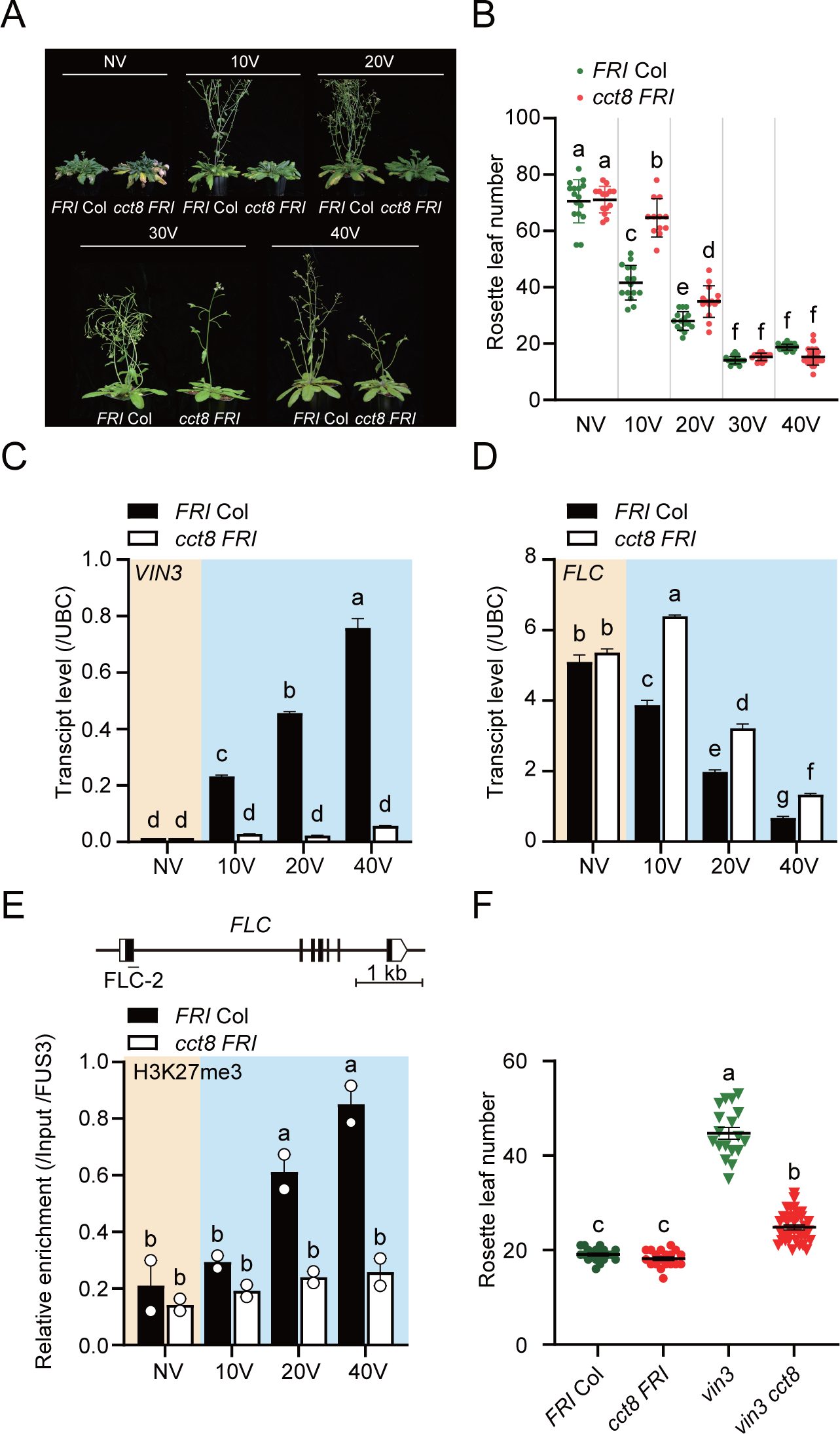
TRiC mediates vernalization response. (A) Representative images of *FRI* Col and *cct8 FRI* after various vernalization periods. (B) Flowering times of *FRI* Col and *cct8 FRI* quantified after various vernalization periods. (C) *VIN3* transcript levels in *FRI* Col and *cct8 FRI* during vernalization time course. (D) *FLC* transcript levels in *FRI* Col and *cct8 FRI* during vernalization. (E) H3K27me3 levels at the *FLC* locus in *FRI* Col and *cct8 FRI* during vernalization. Upper panel: Schematic diagram of the *FLC* gene structure. PCR amplicon for ChIP-qPCR is marked with thin line. Lower panel: ChIP-qPCR results. Relative enrichments of the IP/1% input were normalized to that of *FUS3*. Histograms display mean values ± SEM (n=2 biological replicates, with each biological replicate representing the average of three technical replicates). (F) Flowering times quantified for *FRI* Col, *cct8 FRI*, *vin3 FRI,* and *cct8 vin3 FRI* after 40V. For flowering time analyses, data are presented as means ± SD. Transcript levels were measured using RT-qPCR and are presented as means ± SEM for three biological replicates. Distinct letters indicate a significant difference as determined by two-way ANOVA (B-E) or one-way ANOVA (F) with Tukey’s test.

Compared to the WT, *VIN3* levels in *cct8 FRI* were severely reduced throughout vernalization (Fig. 2C). Similarly, *FLC* levels in *cct8 FRI* were less suppressed compared to the WT during the entire vernalization period (Fig. 2D). Since VIN3 has been reported to recruit PRC2 for the enrichment of H3K27me3 on the *FLC* chromatin (Sung and Amasino 2004), we analyzed the H3K27me3 pattern during vernalization by chromatin immunoprecipitation quantitative PCR (ChIP-qPCR). Although WT showed a gradual increase of H3K27me3 as reported (Yang et al. 2014), *cct8 FRI* showed reduced increase throughout vernalization (Fig. 2E; Supplemental Fig. S3A,B), suggesting that the *VIN3* level in *cct8* is not sufficient to enhance the histone modification, H3K27me3, on the *FLC* chromatin.

Despite reduced *VIN3* level, *cct8 FRI* showed normal acceleration of flowering after 40V compared to WT. Thus, we wondered if *cct8* mutation also affects *VIN3-*independent vernalization response. To address this, we generated *cct8 vin3 FRI* from genetic cross with *vin3-4*, a null allele of *VIN3*. Indeed, *cct8 vin3 FRI* exhibited further acceleration of flowering compared to *vin3 FRI* after 40V although the acceleration was not complete as WT (*FRI* Col) or *cct8 FRI* (Fig. 2F; Supplemental Fig. S3C). Therefore, it suggests that TRiC affects vernalization response both *VIN3* dependently and independently.

Arabidopsis genome contains two more *VIN3* homologs, *VIN3-LIKE 1, 2* (*VIL1, 2*), which have similar structure(Sung et al. 2006; Kim et al. 2010; Kim and Sung 2013; Fiedler et al. 2022; Elsa et al. 2023). Thus, we wondered if any of the VILs compensate for the deficiency of VIN3 in the *cct8* mutant background. When we checked the transcript levels of *VILs* after vernalization, the levels of *VIL1* and *VIL2* were upregulated in *cct8 FRI,* especially in late stages of vernalization (Supplemental Fig. S4). Therefore, the increase of VIL1 and VIL2 may compensate the insufficiency of VIN3 in the *cct8* mutant at late stages of vernalization. Taken together, TRiC is required for the control of *VIN3* and *FLC* expression to achieve proper vernalization response.

### Transcriptome analysis of *cct8* under long-term cold exposure

To reveal the roles of TRiC during prolonged cold, RNA-seq was conducted on NV or 20V WT (*FRI* Col) and *cct8 FRI* plants. The analysis identified a total of 3,682 differentially expressed genes (DEGs) (absolute fold change>1.8199, P<0.05) (Fig. 3A). The gene ontology (GO) enrichment analysis revealed that the DEGs were associated with ‘rhythmic process’, ‘circadian rhythm’, ‘cold acclimation’, ‘cellular transition metal ion homeostasis’ and ‘response to reactive oxygen species’. Particularly, these terms pertained to genes upregulated in WT during cold (WT_20V/NV) and downregulated in *cct8* mutants (20V_mt/WT) (Fig. 3B). To further explore, we identified a total of 94 genes exhibiting expression kinetics similar to *VIN3*; upregulated in WT by 20V but downregulated in *cct8* compared to WT after 20V. We subsequently conducted motif enrichment analysis using the Hypergeometric Optimization of Motif Enrichment (HOMER) software to identify overrepresented cis-elements on their promoters. The analysis revealed enrichment of cis-elements like EE, G-Box, and CRT/DRE in their promoters (Fig. 3C). Notably, EE and G-Box elements are part of the VRE_VIN3_ cis-regulatory element, a vernalization-responsive element, while CRT/DRE has not been detected in the vicinity of the *VIN3* promoter (Kyung et al. 2022). Combined with the GO analyses, these findings suggest that EE, enriched in the promoters of evening-phased clock genes, is potentially involved in the CCT8-mediated *VIN3* regulation. Additionally, our analyses imply that the cold acclimation process, particularly regulated through CRT/DRE elements, is likely to be disrupted in the *cct8* mutant.

**Figure 3.**
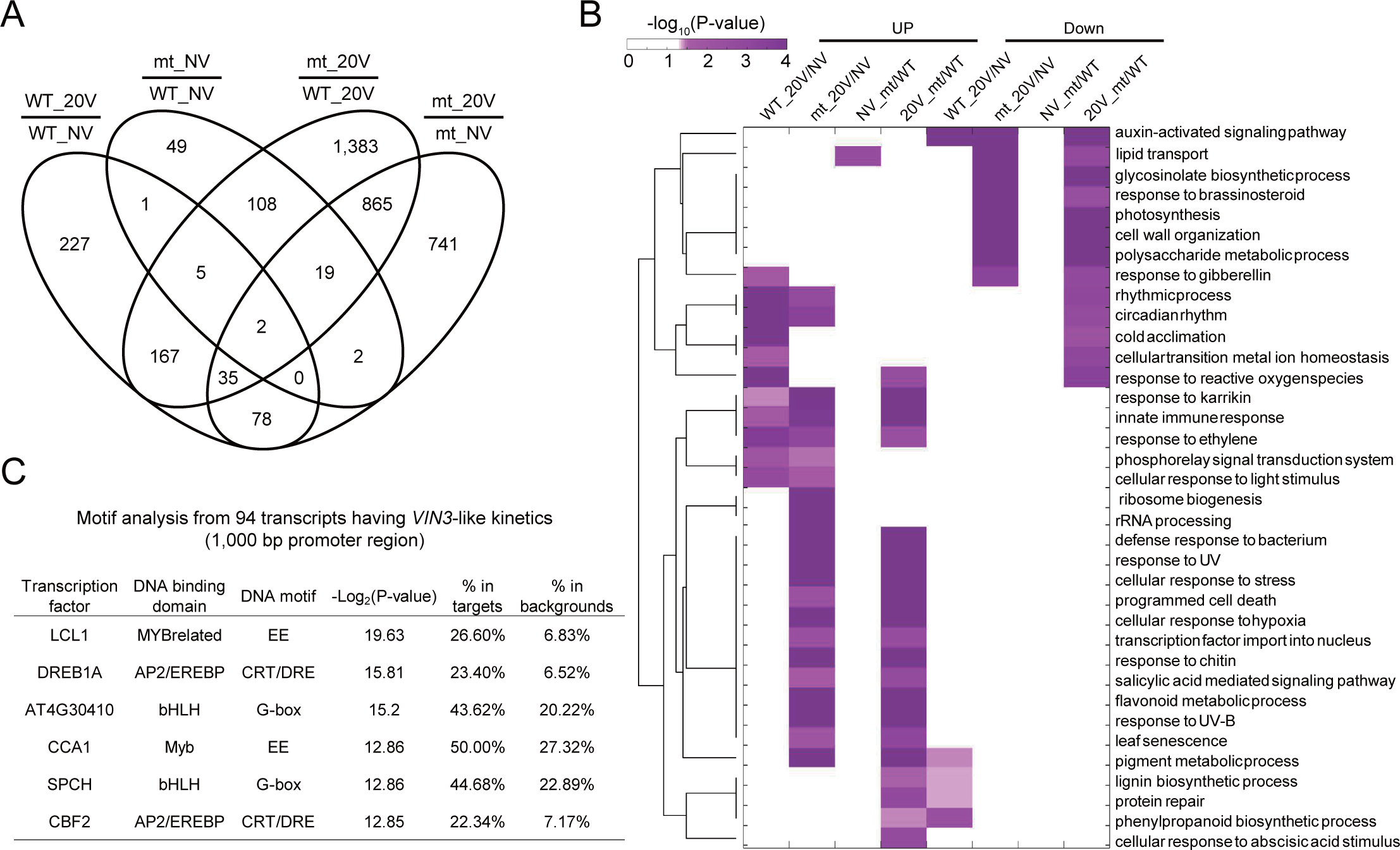
Transcriptome analysis using *cct8* mutants. (A) Venn diagram illustrating differentially expressed genes (DEGs). Legends are as follows: WT (*FRI* Col), mt (*cct8 FRI*), NV (non-vernalized), 20V (20-day vernalized). (B) Gene ontology analysis of DEGs in response to vernalization treatment, presented as a heatmap. The color scale indicates enrichment folds, adjusted for P-value, across different GO terms. The terms WT_20V/NV and mt_20V/NV refer to DEGs in *FRI* Col and *cct8 FRI* due to vernalization, respectively; NV_mt/WT and 20V_mt/WT represent DEGs between WT and mutant under non-vernalized and vernalized conditions, respectively. (C) Result from HOMER DNA-motif enrichment analysis of 94 transcripts displaying expression kinetics akin to *VIN3,* according to the DEG analysis. The motifs column organizes information sequentially, detailing the names of transcription factors, DNA binding domains, and motifs. The table presents the log_2_(p-value) for motif significance, % of target sequences containing the motif, and % of background sequences containing the motif.

### TRiC is required to inhibit anthocyanin accumulation during vernalization

The GO enrichment analysis revealed the association with anthocyanin biosynthesis terms, ‘flavonoid metabolic process’ and ‘pigment metabolic process’ in *cct8* mutants after 20V, coinciding with the observed increase of red pigmentation after 40V (Fig. 1B; Supplemental Fig. S2A). Investigating anthocyanin levels revealed their gradual increase during vernalization in both *cct8-1* and *cct8-5* (Supplemental Fig. S5A,B). Moreover, introducing *CCT8-GFP* transgene restored normal anthocyanin levels in *cct8* mutants (Supplemental Fig. S5B). Consistently, expressions of anthocyanin biosynthesis-related genes (Xu et al. 2017) in the RNA-seq data revealed upregulation of both regulatory genes (*PAP2*, *MYB11*, *MYB12*, and *MYB111*) and structural genes (*CHS*, *F3H*, *PAL1*, *DFR*) in *cct8* mutants after 20V (Supplemental Fig. S6). Furthermore, *HY5* and *HYH*, known upstream regulators of these regulatory and structural genes, were also upregulated in *cct8* mutants after 20V (Supplemental Fig. S6C), supporting their role in promoting anthocyanin biosynthesis, as reported in previous studies (Catalá et al. 2011; Zhang et al. 2011).

Anthocyanin accumulation, typically indicative of stress responses aimed at scavenging reactive oxygen species (ROS) (Falcone Ferreyra et al. 2012), was further evidenced by a substantial increase in H_2_O_2_ levels in vernalized *cct8* mutants (Supplemental Fig. S5C). Collectively, our findings underscore the essential role of chaperonin activity in mitigating exaggerated stress responses triggered by prolonged cold exposure during the vernalization process.

### Circadian rhythm of *VIN3* is regulated by TRiC-RVE module

Our transcriptome analysis revealed an enrichment of “circadian rhythm” and the EE motif among the genes upregulated in the 20V *cct8* mutants (Fig. 3). To confirm the relevance of circadian rhythm, we examined VIN3 levels at 4 hr intervals in both WT and *cct8 FRI* seedlings treated with NV, 20V, and 40V. In non-vernalized conditions, both WT and *cct8 FRI* exhibited a similar diurnal rhythm of *VIN3*, peaking at around 8 h after dawn (zeitgeber time 8; ZT8), albeit with a slightly lower peak in the *cct8 FRI* mutant (Fig. 4A). After vernalization, *VIN3* levels increased, and the peak of *VIN3* rhythm in WT shifted slightly to the daytime (ZT4) after 20V (Fig. 4B). In contrast, *cct8 FRI* displayed not only a severely reduced level but also a loss of rhythmicity after 20V (Fig. 4B,C). Similar effects on the *VIN3* rhythm were also observed in *cct8-1* (Supplemental Fig. S7A), confirming that the *cct8* mutation results in impaired rhythmic expression of *VIN3* during vernalization.

**Figure 4.**
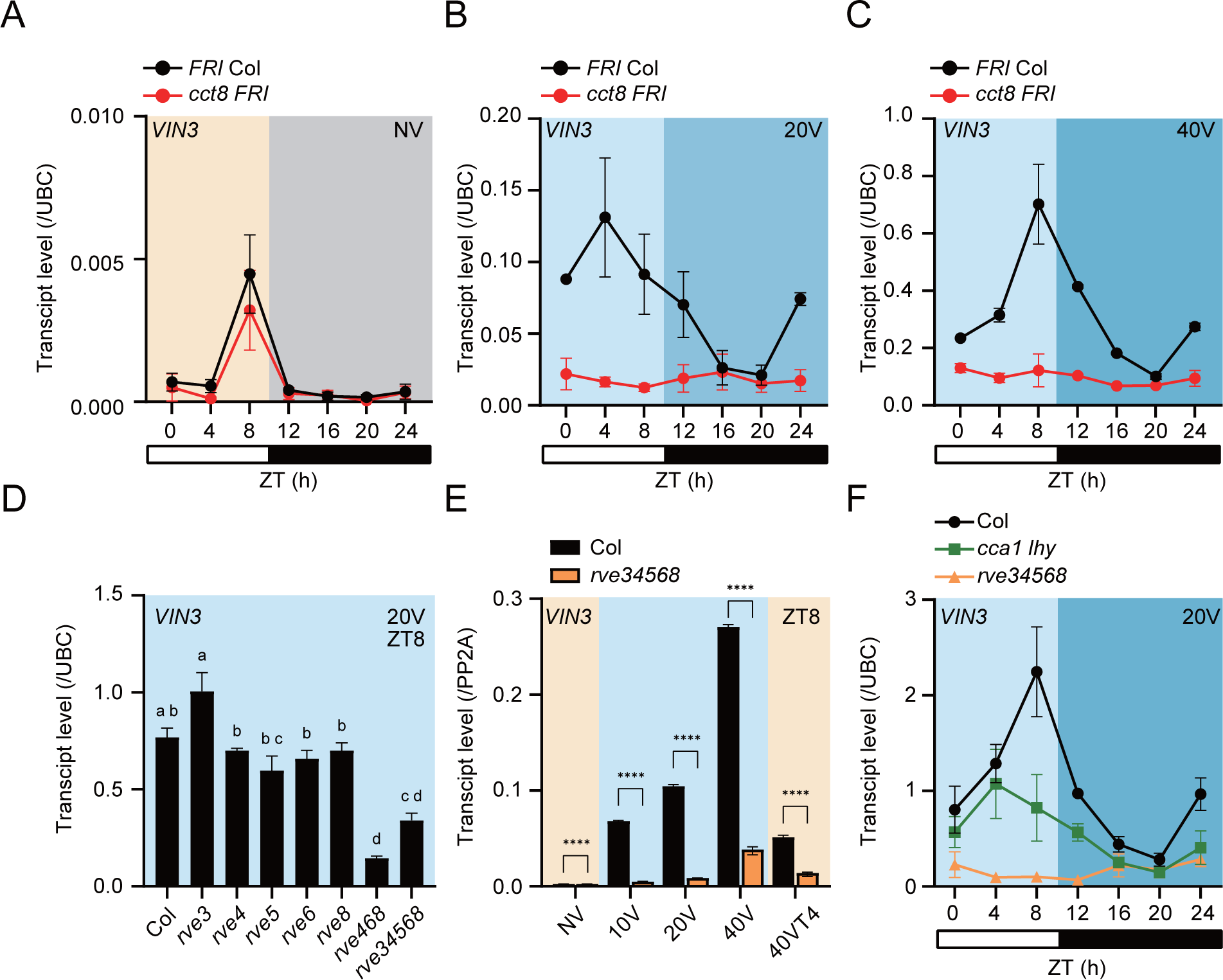
Rhythmic expression of *VIN3* is regulated by TRiC-RVE module. (A-C) Diurnal rhythms of *VIN3* levels in *cct8 FRI* and *FRI* Col before vernalization (A), after 20V (B), and after 40V (C). (D) *VIN3* levels at ZT8 after 20V in Col and various *rve* mutants, including single mutants (*rve3, 4, 5*, *6, 8*), a triple (*rve468*), and a quintuple mutant (*rve34568*). (E) *VIN3* levels in Col and *rve34568* quintuple mutant at ZT8 during vernalization. (F) Diurnal expression of *VIN3* in Col, *cca1 lhy,* and *rve34568* after 20V. Transcript levels were measured using RT-qPCR and are presented as means ± SEM for three biological replicates. White and black boxes represent day and night phases, respectively. Distinct letters denote significant differences (P < 0.05), as determined by one-way ANOVA with Tukey’s test. Asterisks signify significant differences (Student’s t-test; ****P<0.0001).

Consistent with previous reports (Hepworth et al. 2018; Kyung et al. 2022), we observed that the rhythmic expression of *VIN3* was maintained through a free-running experiment conducted under continuous light following 20V (Supplemental Fig. S7B). Then we explored the most sensitive period for *VIN3* induction by cold, *i.e*, gating time of *VIN3.* For this, 10-day-old Col plants grown under short days were transferred to continuous light for 3 days. Then, the plants were subjected to 4 hrs of cold treatment at the indicated ZT time points, followed by the sample collection for *VIN3* level. (Supplemental Fig. S7C). This experimental setup revealed that the most sensitive time for *VIN3* induction by cold is during the daytime, particularly at ZT4 or ZT8, with minimal sensitivity during the subjective night.

Since circadian rhythm in Arabidopsis is regulated by complex networks primarily composed of the evening complex (EC) (Huang and Nusinow 2016), we questioned whether any components of EC or other clock components are responsible for the effect of *cct8* on the arrhythmicity of *VIN3* expression. We found that the diurnal rhythms of *CCA1*, *PRR9*, *PRR7*, *PRR5*, *GI*, and *TOC1* in *cct8 FRI* at 20V were largely intact but generally shifted about 4 hrs towards dusk (Supplemental Fig. S8A). Conversely, the diurnal rhythms and levels of *RVEs* were markedly disrupted by the *cct8* mutation after 20V (Supplemental Fig. S8B). Especially, the transcript levels of *RVE4, RVE5,* and *RVE8* at dawn were significantly reduced in the *cct8 FRI* after 20V (Supplemental Fig. S8B,C). In the case of *RVE8*, the rhythm almost disappeared at the basal level in the *cct8 FRI* after 40V (Supplemental Fig. S8C). Similarly, the *cct8-1* also showed a loss of rhythmicity as well as a strong reduction in *RVE4* and *RVE8* levels compared to WT (Ws-2) (Supplemental Fig. S8D). These results suggest that TRiC is required for the rhythmic expression of *RVEs* during vernalization.

Given the partial redundancy in *RVE* genes (Rawat et al. 2011; Hsu et al. 2013; Gray et al. 2017), we investigated whether mutations in multiple *RVE* genes have a more pronounced effect on *VIN3* expression during vernalization. Among the eight *RVE* family genes, *RVE3, RVE4, RVE5, RVE6,* and *RVE8* are known to have clock functions and act as transcription activators (Farinas and Mas 2011; Shalit-Kaneh et al. 2018). Therefore, we examined *VIN3* levels at ZT8 in single mutants of *RVEs* (*rve3, rve4, rve5, rve6,* and *rve8*), a triple mutant (*rve468*), and a quintuple mutant (*rve34568*) during vernalization (Fig. 4D,E). Every single mutant except *rve3* showed slightly reduced *VIN3* levels, while the triple and quintuple mutants exhibited significantly lower levels compared to WT (Col) (Fig. 4D). Moreover, *rve34568* consistently exhibited a severely reduced *VIN3* level at ZT8 throughout the entire vernalization time course (Fig. 4E).

Finally, we assessed whether the loss of *VIN3* rhythm in the *cct8* mutation could be attributed to circadian clock defects. To this end, we compared the diurnal rhythm of *VIN3* among WT (Col), *cca1 lhy*, and *rve34568* mutants after 20V. As previously reported, WT showed a robust diurnal rhythm, peaking at ZT8 (Kyung et al. 2022), while the *cca1 lhy* double mutant displayed a diminished amplitude, peaking at ZT4 (Fig. 4F). In contrast, the *rve34568* quintuple mutant showed a marked reduction in *VIN3* levels and a complete loss of rhythmicity, mirroring the pattern observed in the *cct8* (Fig. 4B,F).

### RVE-mediated *VIN3* activation is required for vernalization response

We investigated whether RVE proteins directly regulate *VIN3* during vernalization. Gel shift assays with RVE4 and RVE8, expressed as fusion proteins with maltose-binding protein (MBP) (MBP-RVE4, MBP-RVE8) in *E. coli*, demonstrated their direct binding to the VRE_VIN3_ element (Fig. 5A,B; Supplemental Fig. S9). Furthermore, the competition assay using wild-type and mutant versions of the VRE_VIN3_ probe revealed that RVE4 and RVE8 bind specifically to the EE motif in the VRE_VIN3_ element, independent of the G-Box.

**Figure 5.**
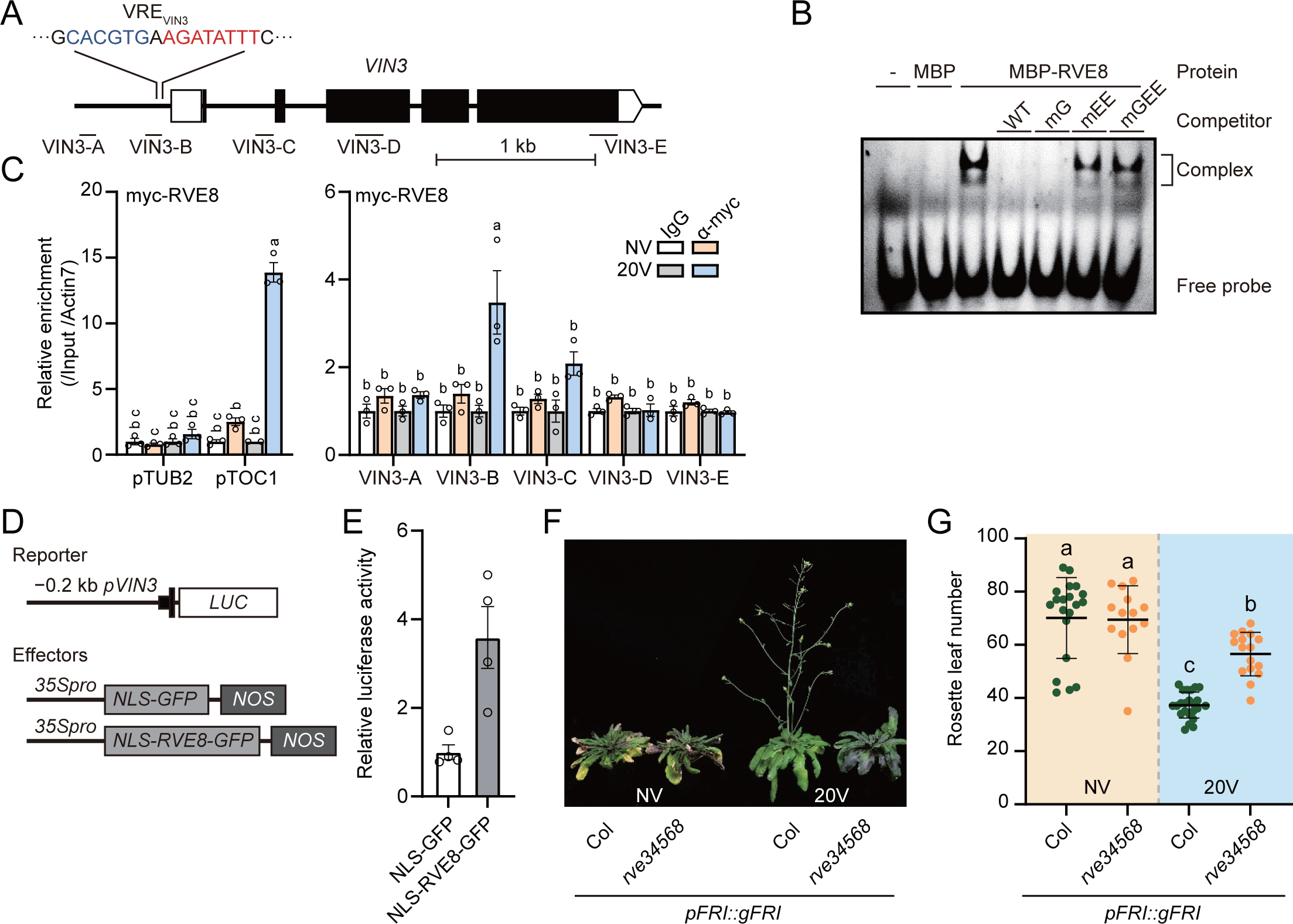
RVE-mediated *VIN3* activation is required for vernalization response. (A) Schematic diagram of the *VIN3* gene structure, illustrating PCR amplicons (VIN3-A to E) used for ChIP analysis as thin lines below the gene structure. The location and sequence of the vernalization responsive *cis*-element in *VIN3* (VRE_VIN3_) are marked with an inverted triangle. (B) Gel shift assay using recombinant RVE8. Unlabeled competitor DNA with a 100× molar excess was introduced to each assay. (C) ChIP-qPCR results showing RVE8 occupancy across the *VIN3* locus in *35S::myc-RVE8* plants, both non-vernalized (NV) and after 20 d of vernalization (20V), harvested at ZT4. Relative enrichments of the IP/1% input were normalized to that of Actin7. The pTUB2 and pTOC1 served as negative and positive controls, respectively. Data are presented as means ± SEM for three biological replicates, with individual data points shown as open circles. (D) Diagram of reporter and effector constructs for luciferase assay. The reporter vector includes a −0.2 kb promoter, 5′-UTR, and the 1^st^ exon of *VIN3,* fused with the *luciferase* (*LUC*) gene. (E) Luciferase activity in co-transfected *Arabidopsis* mesophyll protoplasts. Relative luciferase activity was normalized to the NLS-GFP control. Data are shown as means ± SEM for four biological replicates, with individual data points as open circles. (F) Representative photographs of primary *Arabidopsis* transgenic lines carrying the genomic *FRI* transgene in either Col or *rve34568,* with or without 20V. (G) Flowering time quantification of transgenic lines, expressed as means ± SD. For (C) and (G), Distinct letters indicate a significant difference (P < 0.05) as determined by two-way ANOVA with Tukey’s test. For (E), asterisks signify significant differences (Student’s t-test; *P<0.05).

*In vivo* binding of RVEs to the *VIN3* promoter was confirmed by ChIP-qPCR using *35S::myc-RVE8* transgenic plants (Fig. 5A,C). The myc-RVE8 was significantly enriched in the promoter region of *VIN3* near the EE motifs only after 20V, indicating direct binding of RVE8 to the *VIN3* promoter. This finding is consistent with a previous report demonstrating that RVE8 is translocated to the nucleus under cold stress to regulate target genes (Kidokoro et al. 2021).

The transcriptional activation of *VIN3* by RVE8 was further confirmed by an Arabidopsis protoplast transfection assay. To facilitate translocation to the nucleus, RVE8 was fused with Nuclear Localization Signal (NLS). The transient expression of the NLS-RVE8-GFP fusion protein significantly increased the activity of the reporter luciferase controlled by the *VIN3* promoter, compared to the control NLS-GFP (Fig. 5D,E). Taken together, our results strongly suggest that the RVE8 protein, translocated to the nucleus, directly activates *VIN3* expression by binding to the EE motif in the *VIN3* promoter during vernalization.

Finally, we investigated whether *RVEs* are necessary for the proper vernalization response. To address this, we introduced *pFRI::gFRI* into WT (Col) and *rve34568* quintuple mutant via *Agrobacterium*-mediated transformation, enabling vernalization sensitive response (Michaels and Amasino 1999). The first-generation transgenic plants were compared for flowering time after NV or 20V. While the flowering time of *pFRI::gFRI* and *pFRI::gFRI rve34568* without vernalization was similar, *pFRI::gFRI rve34568* plants exhibited significantly delayed flowering compared to *pFRI::gFRI* plants after 20V (Fig. 5F,G). These results support the notion that *RVEs* are indispensable for the proper vernalization response in flowering.

### Overexpression of *RVE8* rescues *rve34568* but fails to rescue *cct8* for *VIN3* activation

To assess the role of *RVE8* in *VIN3* activation, we introduced *35S::myc-RVE8* into the *rve34568* mutant and compared *VIN3* levels among WT (Col), *rve34568,* and *35S::myc-RVE8 rve34568* transgenic plants after 20V at ZT4. The introduction of *35S::myc-RVE8* significantly increased *VIN3* levels in *rve34568* (Fig. 6A,B), indicating that *RVE8* overexpression rescues the defect in the *rve34568* mutant. However, introducing *35S::myc-RVE8* into the *cct8* mutant could not restore *VIN3* activation throughout the vernalization time course (Fig. 6C).

**Figure 6.**
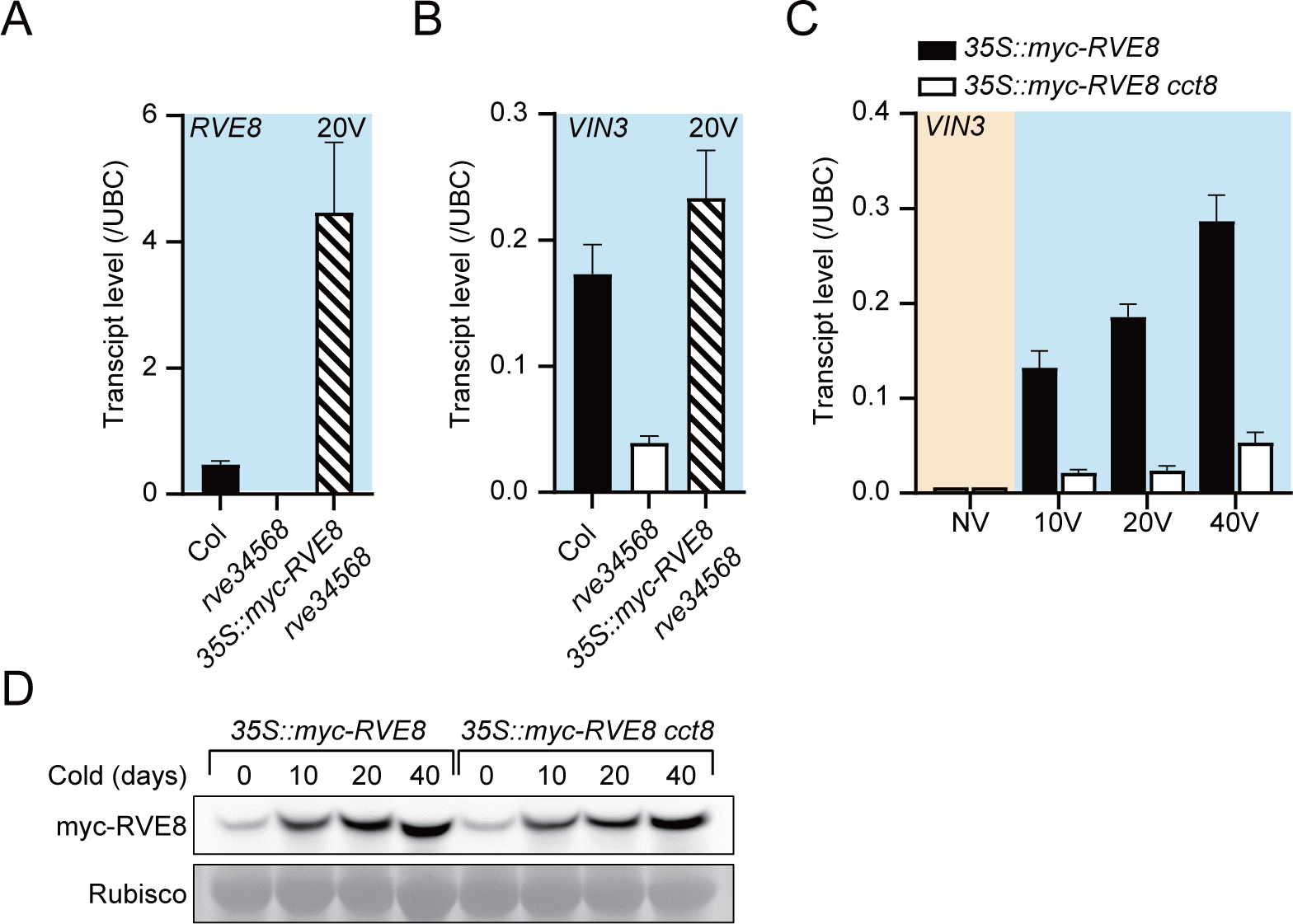
Overexpression of *RVE8* rescues *rve34568* but fails to rescue *cct8* for *VIN3* activation (A-B) Expression levels of *RVE8*. (A) and *VIN3* (B) in the seedlings of WT (Col), *rve34568*, and *35S::myc-RVE8 rve34568* after 20V. (C) *VIN3* transcript levels in the seedlings of *35S::myc-RVE8* and *35S::myc-RVE8 cct8* during vernalization. (D) Immunoblot analysis showing abundance of the myc-RVE8 protein in *35S::myc-RVE8* and *35S::myc-RVE8 cct8* during vernalization. Rubisco was used as loading control. Transcript levels were measured using RT-qPCR and presented as means ± SEM for three biological replicates.

Since *CCT8* is a component of chaperonin involved in protein stability, the failure of *VIN3* activation in *35S::myc-RVE8 cct8* may be due to the degradation of RVE8 protein in the absence of TRiC. To address this, we compared the RVE8 protein levels in *35S::myc-RVE8 cct8* and *35S::myc-RVE8* during vernalization (Fig. 6D). The results showed that the RVE8 protein level in the mutant (*35S::myc-RVE8 cct8*) is comparable to that in the WT (*35S::myc-RVE8*). Collectively, our results suggest that RVEs are necessary for proper *VIN3* activation, but the ectopic expression of *RVE8* alone is not sufficient to restore vernalization-induced *VIN3* activation in *cct8*. This indicates that the *cct8* mutation causes defects not only in the circadian oscillator, RVEs, but also in other factor(s), likely G-box binding factor(s) (Kyung et al. 2022).

### TRiC-RVE module mediates cold acclimation under long-term winter cold

Our transcriptome analysis revealed an enrichment of “cold acclimation”, and a enrichment of the CRT/DRE motif in *VIN3*-like kinetics genes (Fig. 3). This led us to explore whether TRiC is also essential for cold acclimation. Comparison of freezing tolerance between WT (*FRI* Col) and *cct8 FRI* after 20V revealed a significant decrease in tolerance in *cct8 FRI* (Fig. 7A,B), suggesting that TRiC plays a role in cold acclimation after long-term cold exposure. However, this result prompts the question of whether TRiC is involved in the previously observed gradual increase of *CBFs* during long-term cold (Jeon et al. 2023).

**Figure 7.**
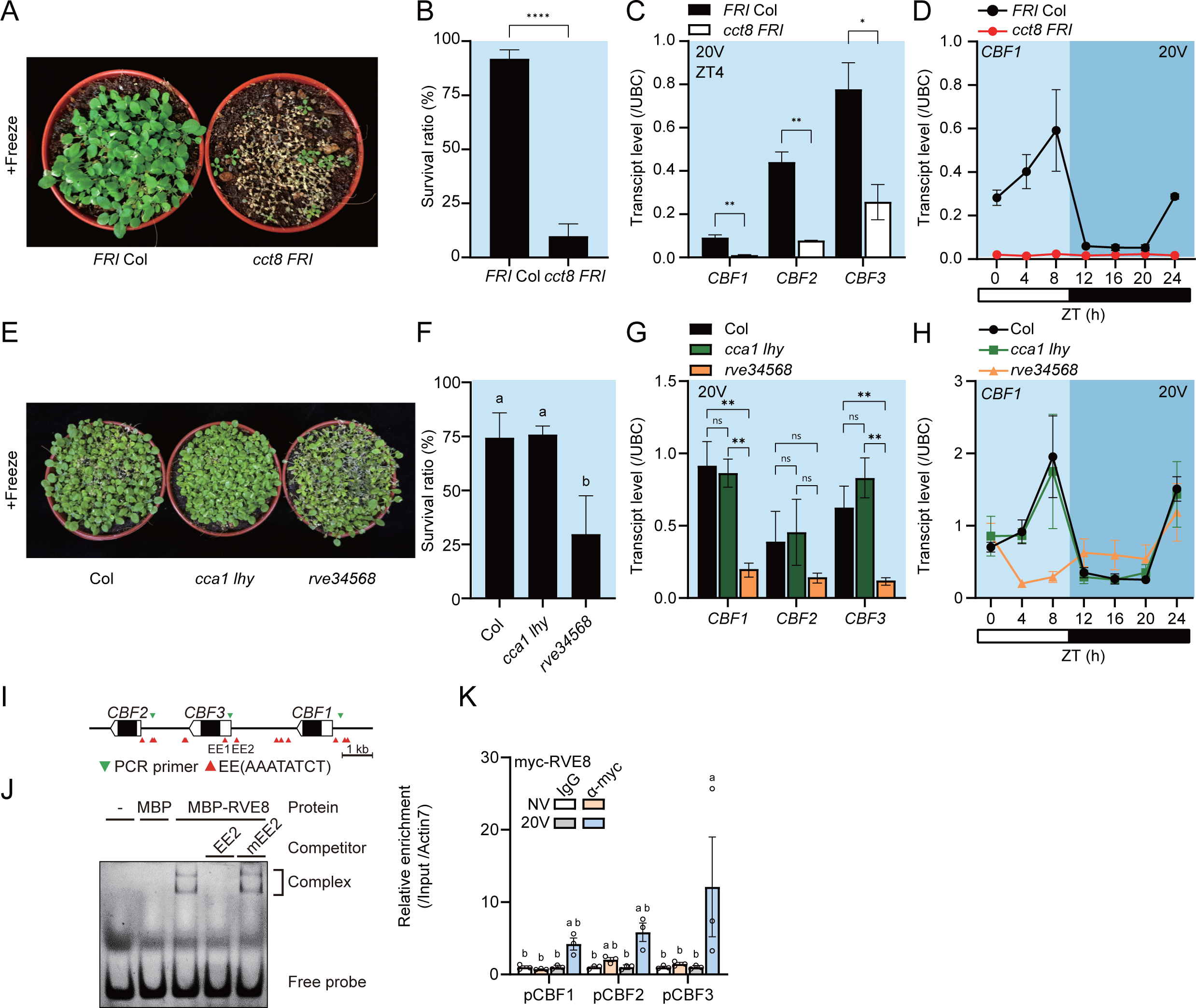
TRiC-RVE module facilitates long-term cold specific acclimation. (A-B) Freezing tolerance of the seedlings of *FRI* Col and *cct8 FRI* after 20V, exposed to −10°C for 3 h. (A) Representative images and (B) survival ratios after 7 days of recovery at room temperature. (C) Transcript levels of *CBFs* in the seedlings of WT and *cct8* after 20V at ZT4. (D) Diurnal gene expression of *CBF3* in *FRI* Col or *cct8 FRI* after 20V. (E-F) Freezing tolerance in the seedlings of Col, *cca1 lhy* and *rve34568* after 20V, subjected to −10°C for 3 h. (E) Representative images and (F) survival ratios after 7 days of recovery at room temperature. (G) Transcript levels of *CBFs* in the seedlings of WT and the mutant after 20V at ZT4. (H) Diurnal *CBF1* expression in Col, *cca1 lhy* or *rve34568* after 20V. (I) Schematic of the genomic region encompassing *CBFs*, highlighting EE sites (red arrowheads) and ChIP-qPCR amplicons (green arrowheads). (J) Gel shift assay using recombinant RVE8 and Cy5-labeled EE2 probes, including competitor DNAs (100× molar excess) with WT (EE2) or mutated EE2 (mEE2) (K) RVE8 occupancy at *CBF* gene promoters via ChIP-qPCR in *35S::myc-RVE8* plants, NV or after 20V, at ZT4. Relative enrichments of the IP/1% input were normalized to that of Actin7. Data are presented as means ± SEM for three biological replicates, with individual data points shown as open circles. Transcript levels were measured using RT-qPCR and presented as means ± SEM for three biological replicates. For (A) and (E), data are presented as means ± SD from three biological replicates. For (B) and (C), asterisks signify significant differences (Student’s t-test; *P<0.05, **P<0.01, ****P<0.0001). For (F) and (G), distinct letters indicate a significant difference (P < 0.05) based on one-way ANOVA with Tukey’s test.

We compared transcript levels of *CBF1*, *CBF2*, *CBF3*, and *RD29a* in WT and *cct8 FRI* during short-term (0, 1, 3, 6, 12, 24 h) and long-term (0, 20, 40 d) cold treatment. Transcript levels of all the *CBFs* and *RD29a* during short-term cold treatment were similar in WT and *cct8 FRI* (Supplemental Fig. S10A), peaking after 3 hours of cold exposure and gradually decreasing to basal levels, consistent with a previously reported pattern (Gilmour et al. 1998). However, during long-term cold treatment, while transcript levels of *CBFs* in WT gradually re-increased after 20V, those in *cct8 FRI* were severely reduced and reached basal levels after 40V (Fig. 7C; Supplemental Fig. S10B). Similarly, WT showed a strong increase in *RD29a* after 20V but *cct8 FRI* barely induced (Supplemental Fig. S10B). These results confirm that there are two waves of *CBFs* expression, one for short-term and another for long-term cold exposure, as previously reported (Jeon et al. 2023), and TRiC is involved only in the long-term cold response. Furthermore, these findings strongly suggest that the TRiC-mediated cold acclimation process is critical for the survival of plants during long-term winter cold.

As *RVE4* and *RVE8* were reported to activate *CBFs* under cold stress (Kidokoro et al. 2021), we investigated whether the TRiC-RVE module is associated with the reduced cold acclimation capacity in *cct8*. We compared the rhythmic expression of *CBFs* and *RD29a* in WT and *cct8 FRI* during vernalization treatment. As expected, the mutant showed reduced, and arrhythmic expression of *CBFs* and *RD29a* after 20V (Fig. 7D; Supplemental Fig. S10C). Moreover, freezing tolerance experiments conducted after 20V revealed a significantly reduced survival rate of the *rve34568* quintuple mutant, while the survival rate of *cca1 lhy* was comparable to that of WT (Fig. 7E,F), indicating that the cold acclimation observed after long-term cold exposure is specific to *RVEs*. Consistently, *rve34568* mutants showed arrhythmic and reduced levels of *CBF1, CBF3,* and *RD29a*, while *cca1 lhy* mutants maintained rhythms similar to WT (Fig. 7G,H; Supplemental Fig. S10D). In case of *CBF2*, the effect of the mutations was relatively minor in both *rve34568* and *cca1 lhy* (Supplemental Fig. S10D). Notably, *CCA1* rhythm diminished in the *rve34568* mutant, but *RVE8* rhythm remained stable in *cca1 lhy* mutants after 20V (Supplemental Fig. S10E). This indicates that *CCA1*’s rhythm depends on *RVEs* and underscores the role of RVEs in modulating cold acclimation through *CBFs* during winter cold.

We then investigated whether *RVE8* overexpression could rescue the defect of cold acclimation observed in the *cct8* mutant. Similar to the impaired *VIN3* activation after vernalization in *35S::myc-RVE8 cct8*, *35S::myc-RVE8* failed to induce the expression of *CBFs* and *RD29a* after vernalization in *cct8* mutants, although it successfully rescued such failure in the *rve34568* mutant (Supplemental Fig. S11A,B). Furthermore, *35S::myc-RVE8* did not rescue cold acclimation defect in *cct8* mutants (Supplemental Fig. S11C). These findings suggest that RVEs are not sufficient but additional factor(s) are necessary for cold acclimation after winter cold, similar to the vernalization response.

To confirm the direct regulation of *CBFs* by RVEs, we conducted gel shift assay with MBP-RVE8 and MBP-RVE4 purified from *E. coli.* The results showed that both recombinant proteins bind to DNA probes of EE1_CBF3_ or EE2_CBF3_ with flanking sequences, whereas they failed to interact with the mutated version of the probes (Fig. 7I,J; Supplemental Fig. S12). The *in vivo* interaction between RVE8 and *CBF* promoters was investigated by ChIP-qPCR. Notably, myc-RVE8 was found to be significantly enriched after 20V but not before cold treatment (Fig. 7K), consistent with previous reports indicating that RVE8 activates *CBFs* under cold stress (Kidokoro et al. 2021). Taken together, our results suggest that the TRiC-RVE module also mediates cold acclimation via RVEs, by directly activating the transcription of *CBFs* during winter cold.

## Discussion

Plants, notably Arabidopsis, endure prolonged winter cold, yet the mechanisms enabling their survival under such harsh conditions have remained elusive (Larran et al. 2023). Additionally, while the biochemical functions of the chaperonin have been extensively studied, its biological significance has received comparatively less attention. In our study, we elucidate the indispensable role of the TRiC complex in plant fitness during prolonged cold exposure, operating through the regulation of key components such as the circadian clock, RVEs, VIN3, and CBFs, which are pivotal for the vernalization response and cold acclimation.

Beyond previously reported functions of Arabidopsis TRiC, our study highlights that the TRiC is crucial for orchestrating responses to chronic environmental changes, exemplified by winter cold. Our data highlight a role for TRiC as an upstream component of RVEs, key clock transcription factors. TRiC is required for the rhythmic expression of *RVEs* under prolonged cold, but not for the other clock genes (Supplemental Fig. S8), suggesting the specificity of TRiC-mediated winter cold responses. The regulation of *RVEs* has been shown to underlie two distinct processes triggered by prolonged cold: vernalization and cold acclimation. Given that both *cct8* and *rve34568* mutants showed delayed flowering and compromised cold acclimation after vernalization treatment (Figs. 2A,B and 4F), it is evident that TRiC-RVE module impacts both of the winter responses. Nevertheless, the precise mechanism underlying TRiC-mediated protein folding and transcriptional regulation of RVE genes remains unclear. The *cct8* mutant exhibits temperature-dependent pleiotropy such as retarded growth and pigment accumulation (Fig. 1B). The upregulation of *HY5*/*HYH* in the *cct8* mutant, which are key players in light signaling and anthocyanin biosynthesis, suggests that TRiC also plays a role in tempering the excessive stress response triggered by prolonged cold (Supplemental Fig. S5). These findings provide valuable insights into the link between the observed phenotype and the processes of winter adaptation, vernalization and cold acclimation.

Our findings establish that RVE8 directly binds to *VIN3* and *CBFs*, essential regulators of vernalization and cold acclimation respectively, thus activating their transcription and orchestrating proper responses to prolonged cold (Figs. 5 and 7). Our findings emphasize the importance of *RVEs* in mediating prolonged cold acclimation, whereas previous studies have focused on the circadian clock’s regulation of cold induction of *CBFs* for short-term cold acclimation (Dong et al. 2011; Keily et al. 2013; Liu et al. 2013). The perception of long-term cold by RVEs is shared by both vernalization and cold acclimation processes. This shared regulatory mechanism that shapes the diurnal expression pattern of *VIN3* and *CBFs* suggests that the circadian clock plays a crucial role in the signaling/perception of long-term cold exposure, facilitating differential responses to day and night temperatures. The ability to differentiate between winter and sudden temperature drops may align with the evolutionary need for resource allocation in preparing for long-term winter cold. However, the factors leading to the increased expression of *VIN3* and *CBFs* throughout the duration of cold exposure remain to be addressed.

The *VIN3* promoter contains VRE_VIN3_, a region crucial for *VIN3* activation, comprising G-box and EE DNA motifs. Previous studies have demonstrated that mutations in either EE or G-box result in defects of partial *VIN3* activation, while mutations in both elements abolish the response, indicating the necessity of both EE-binding and G-Box-binding transcription factors for proper *VIN3* activation (Kyung et al. 2022). Notably, the G-box motif is frequently found in the promoters of morning-phased genes regulated by CCA1 (Nagel et al. 2015), implying potential interaction between EE-binding transcription factors and unidentified G-box binding factor(s), thereby synergistically activating their target genes. In our study, we identified the circadian clock transcription factor RVE as the binding factor for EE motifs on the promoters of *VIN3* and *CBFs*, which contain both G-box and EE motifs. However, the specific identity of the transcription factor(s) binding to the G-box motif remains unknown. Further studies are warranted to elucidate these transcription factors and unravel the intricate regulatory mechanisms underlying prolonged cold conditions.

*FLC* is a major regulator of flowering time in *Arabidopsis,* but it is not the sole factor involved in flowering regulation. Arabidopsis flowering is governed by intricate pathways, encompassing both *FLC*-dependent and *FLC*-independent mechanisms. The *cct8 vin3 FRI* mutant exhibits accelerated flowering compared to the *vin3 FRI* after vernalization treatment, albeit *FLC* levels are statistically not different (Fig. 2F; Supplemental Fig. S3C). It suggests that TRiC is also involved in the *FLC*-independent flowering pathway. In addition, vernalization primarily regulates *FLC* level by the epigenetic suppression through both *VIN3*-dependent and *VIN3*-independent pathways, as previously reported (Hepworth et al. 2018). Although the *rve34568* shows a defect of vernalization response (Fig. 5F), *cct8* mutants, characterized by an arrhythmic *RVE* expression, exhibit the defective response only in the early phase of vernalization (Fig. 2A–C). Considering the role of TRiC as a general protein folding machinery, the differential response between *cct8* and *rve34568* mutants implies the involvement of TRiC in *VIN3-*independent *FLC* repression. Thus, TRiC is broadly implicated in both *VIN3*-dependent/independent and *FLC*-dependent/independent pathways.

During vernalization, rhythmic expression of *VIN3* was disrupted in *cct8* mutants, yet *VIN3* levels at nighttime were similarly low in both wild type and *cct8* mutants (Fig. 4B), albeit higher than those observed in non-vernalized plants. This suggests that the basal level of *VIN3* is elevated with vernalization, and RVE-mediated transcriptional activation contributes to rhythmic *VIN3* expression. Considering that EE and G-box govern the activity of the *VIN3* promoter during prolonged cold exposure (Kyung et al. 2022), it is likely that the G-box is responsible for elevating the basal expression level of *VIN3*. Additionally, as *Arabidopsis* can be vernalized even in the dark (Chandler and Dean 1994), during which clock gene expression becomes arrhythmic (Millar et al. 1995), EE-mediated regulation likely contributes to rhythmic *VIN3* expression by integrating environmental signals through circadian oscillators for optimal developmental transition.

The upregulation of cold-responsive genes in a day-specific manner under prolonged cold exposure suggests a mechanism for differential response to day and night temperatures. Given that nights are typically colder than days due to the absence of sunlight, plants might interpret cold days, despite the presence of sunlight, as indicative of the onset of long-term cold, *i.e*., winter. This ability to distinguish between temporal temperature drops and prolonged cold may have evolved because preparing for long-term winter cold demands significant resources. Our interpretation aligns with previous studies showing that *VIN3* expression increases during constant cold without temperature fluctuation, as opposed to temperatures that fluctuate around the same average but including relatively higher temperature at daytime (Antoniou-Kourounioti et al. 2018; Hepworth et al. 2018). Although it remains uncertain whether *CBFs* are suppressed by warm spikes during long-term cold, the similar diurnal oscillation patterns of *VIN3* and *CBFs* suggest that the amplified expression by RVE8 during the daytime serves as a positive mechanism for responding to prolonged cold.

In summary, our study provides novel insights into the intricate interplay between the TRiC, the circadian clock, and key transcription factors (RVEs), shedding light on their collective roles in orchestrating plant responses to prolonged winter cold. The shared regulatory mechanism through RVEs in vernalization and cold acclimation processes unveils a previously unrecognized connection between these two critical environmental adaptations.

## Materials and Methods

### Plant Materials and Growth Conditions

All Arabidopsis thaliana lines used in this study were of the Columbia (Col-0) background, except for the Landsberg *erecta* (L*er*) ecotype, which was used to generate a mapping population for map-based gene cloning. Additionally, the Wassilewskija (Ws-2) ecotype was employed to examine the *cct8-1* mutant. The wild type, Col:*FRI^Sf2^* (*FRI* Col) has been previously described (Lee et al. 1994). The *vin3-4*, *rve34568* and *cca1 lhy* mutants have been previously described (Gray et al. 2017; Kyung et al. 2022).

To construct *35S::myc-RVE8*, RVE8-coding sequences were amplified and fused in-frame with *myc-pBA* vector. The *35S::myc-RVE8 cct8* transgenic lines were produced by crossing *35S::myc-RVE8* with the *cct8-1* mutant. For creating *35S::myc-RVE8 rve34568* lines, coding sequence of *myc-RVE8* was amplified using PCR and cloned into *pPZP211* plasmid containing the *35Spro* and NOS terminator, then transformed into the *rve34568* mutant. The *pFRI::gFRI* construct was generated by amplifying genomic sequences that include 1 kb upstream of the promoter and the FRI-coding sequence from *FRI^Sf2^*, which were then inserted into the *pPZP211* vector. These constructs were introduced into the indicated lines using the *Agrobacterium tumefaciens*-mediated *Arabidopsis* floral dip method (Clough and Bent 1998).

Plants were grown under 16 h/8 h light/dark cycle (long day) or 8 h/16 h light/dark cycle (short day) (22 ℃/20 ℃) in a controlled growth room lit by cool white fluorescent lights (125 μmol m^-2^ sec^-1^). Vernalization treatments were done as previously described (Kim et al. 2010). Non-vernalized seedlings were grown for 11 d. For 10V, 20V, 40V treatments, seedlings were grown under short days for 10, 9, 7 d, respectively, after germination and then transferred to a vernalization chamber at 4 ℃. Following vernalization treatment, seedlings were either collected or transplanted into soil. Flowering time was measured by counting the number of rosette leaves at the opening of the first flower using at least 12 plants.

### EMS mutagenesis and positional cloning

EMS mutagenesis was conducted as previously described (Kim et al. 2006). For the positional cloning of the causative gene of *P161*, F2 progenies were obtained by crossing *P161* to L*er*. To identify the gene responsible for the phenotype observed in *P161*, F2 progenies were generated by crossing *P161* with the L*er* ecotype. The mapping procedure utilized 94 GUS-inactive F2 plants and molecular markers, as detailed in previous studies (Lukowitz et al. 2000; Hou et al. 2010). After rough mapping, the genomes of *P161* and the parental −0.2 kb *pVIN3_U_I::GUS* lines were sequenced and compared by illumina Hiseq4000 platform (Illumina, San Diego, CA, USA) to find mutant-specific SNPs in *P161* using LabGenomics services.

### Histochemical Staining

GUS staining was carried out using standard methods as previously described (Shin et al. 2018). Seedlings were grown at room temperature for 10 d, followed by the exposure to vernalization (8 h/16 h light/dark cycle at 4°C). For 3,3′-diaminobenzidine (DAB) staining, Arabidopsis seedlings were submerged in a DAB solution (1 mg/mL DAB, pH 3.8) for 16 hours. Following staining, the seedlings were destained by immersing them in 100% ethanol for 3 hours. Photographs were taken using a Dimis-M USB digital-microscope (Siwon Optical Technology).

### Quantitative PCR

For real-time quantitative PCR analysis, total RNA was extracted using TRIzol solution (Sigma-Aldrich). Four micrograms of total RNA were treated with recombinant DNaseI (TaKaRa) to eliminate genomic DNA. cDNA synthesis was then performed on the DNase I-treated RNA using reverse transcriptase (Thermo Scientific, EP0441) and oligo(dT) primers. Quantitative PCR was conducted using the 2× SYBR Green SuperMix (Bio-Rad) on a CFX96 real-time PCR detection system. The relative transcript levels of each gene were normalized to those of *UBC* or *PP2A*, serving as the reference gene.

### Chromatin Immunoprecipitation

Approximately 4 g of whole *Arabidopsis* seedlings were collected, cross-linked with 1% (v/v) formaldehyde for 10 min, and then quenched with 0.125 M glycine for 5 min under vacuum. The seedlings were washed with distilled water, immediately frozen in liquid nitrogen, and ground into a fine powder. The powder was then resuspended in Nuclei Isolation Buffer (1 M hexylene glycol, 20 mM PIPES-KOH (pH 7.6), 10 mM MgCl_2_, 15 mM NaCl, 1 mm EGTA, 1 mM PMSF, and complete protease inhibitor mixture tablets (Roche)). *Arabidopsis* nuclei were isolated by centrifugation, lysed by Nuclei Lysis Buffer (50 mM Tris-HCl (pH 7.4), 150 mM NaCl, 1% Triton X-100, 1% SDS), and sonicated with a Branson sonifier to shear the DNA into fragments. The resulting sheared chromatin solution was diluted 10-fold with a ChIP Dilution Buffer (50 mM Tris-HCl (pH 7.4), 150 mM NaCl, 1% Triton X-100, 1 mM EDTA). This diluted chromatin was then incubated overnight at 4°C with Protein A/G PLUS-Agarose beads (Santa Cruz Biotechnology; sc-2003) and antibodies: either anti-trimethyl-Histone H3 (Millipore; 07-449), anti-myc (Santa Cruz Biotechnology; sc-40), or normal mouse IgG1 (Santa Cruz Biotechnology; sc-3877) as a control. The beads were washed 4 times with ChIP dilution buffer, and DNA was extracted using Chelex 100 resin according to the manufacturer’s instruction. qPCR analysis was then performed using 1% input and immunoprecipitated DNA.

### Transcriptome analysis

Total RNA was isolated from *Arabidopsis* plants as described above. Bioanalyzer RNA Pico 6000 chip was applied to qualify sample availability. An RNA-Seq library was constructed using the TruSeq RNA Sample Preparation kit (Illumina), following the manufacturer’s instructions. Sequencing of the RNA-Seq library was carried out to produce paired-end reads (2 × 100 bp) using the Illumina Novaseq system (Illumina), with sequencing services provided by Macrogen. The genes with fragments per kilobase of exon model per million reads mapped (FPKM) values larger than 0 in at least one of the samples for each condition was defined as expressed genes. The read counts of expressed genes were normalized using the TMM normalization method in the edgeR package ver 3.6 (Robinson et al. 2010). Subsequently, the normalized read counts were log2-transformed and further normalized using quantile normalization (Robinson et al. 2010). Afterward, T-statistic values and log2-fold-changes for each gene in comparisons were computed. Empirical null distributions of the T-statistic values and log2-fold-changes were estimated by performing random permutations of all samples 1,000 times. Instead of conventional fold-change cutoffs, significance thresholds using the estimated empirical distributions of log2-fold-change values were established. An average of top 2.5% and 97.5% absolute log2-fold-change values (1.8199) was selected. Finally, DEGs were identified as the genes that had t-test p-values < 0.05, and absolute log2-fold-changes > 1.8199. The enrichment analysis of GOBPs for up- and downregulated gene sets in each comparison was performed using DAVID software ver. 6.8 (Huang et al. 2009). GOBPs with Benjamini-Hochberg p-value < 0.05 were defined as the processes enriched in the gene set. Motif enrichment analysis was performed on the promoters of 94 genes exhibiting *VIN3*-like kinetics (WT_20V/NV fold-change > 1.8199, mt_20V/NV < - 1.8199, p-value < 0.05) in the gene expression profiles. HOMER (Hypergeometric Optimization of Motif EnRichment) ver. 4.11 (Heinz et al. 2010) was employed, with the TAIR10 reference genome and a motif region size of 1000 bp provided. The TAIR10 reference genome sequence (GCF_000001735.4) was obtained from the RefSeq database. Total of 506 motifs were discovered through de novo motif finding, including 60 known motifs significantly enriched in cis-regulatory elements of genes exhibiting *VIN3*-like kinetics (Benjamini-Hochberg p-value < 0.01).

### Quantification of anthocyanin contents

Anthocyanin contents were measured using previously described methods (Mita et al. 1997) Briefly, homogenized seedlings were incubated with methanol-HCl (1%, v/v) at 4°C for 24 h in the dark. After centrifugation at 16,000 g for 15 min, the supernatant was collected, and absorbance was measured at 657 nm and 530 nm. The anthocyanin level was calculated with the formula: (A530-0.25*A657)/fresh weight (g).

### Immunoblotting

For immunoblot assay, total proteins were prepared from 100 mg of harvested samples in protein extraction buffer (50 mM Tris-Cl pH 7.5, 150 mM NaCl, 10 mM MgCl_2_, 1 mM EDTA, 1% Triton X-100, 1 mM PMSF, 1 mM DTT, 1× complete Mini, and EDTA-free protease inhibitor cocktail (Roche). Total proteins were separated by SDS-PAGE. The proteins were transferred to PVDF membranes (Cytiva) and probed with anti-α TUB (Santa Cruz Biotechnology; sc-23948, 1:10,000 dilution) or anti-myc (Santa Cruz Biotechnology; sc-40, 1:10,000 dilution) antibodies overnight at 4 ℃. The samples were then probed with horseradish peroxidase-conjugated anti-mouse IgG (Cell Signaling; #7076, 1:10,000 dilution) antibodies at room temperature. The signals were detected using ImageQuant LAS 4000 (Cytiva) with WesternBright^TM^ Sirius ECL solution (Advansta).

### Preparation of fusion protein and gel shift assay

Coding sequence encoding RVE4 and RVE8 was fused to *pMAL-c2* vector. The MBP, MBP-RVE4 and MBP-RVE8 proteins were expressed in *Escherichia coli* BL21 strain according to the manufacturer’s instructions using the pMAL Protein Fusion and Purification System (New England Biolabs; #E8200) and purified using MBPtrap HP column (Cytiva) attached to ÄKTA FPLC system (Cytiva). The Cy5-labeled probes and unlabeled competitors were generated by annealing 40 bp-length oligonucleotides (Table S1). 5 μM of purified proteins and 100 nM of Cy5-labeled probe were incubated at room temperature in binding buffer (10 mM Tris-HCl (pH 7.5), 50 mM NaCl, 1 mM EDTA, 5% glycerol and 5 mM DTT). For competition assay, 100× molar excess of each competitor was added to the reaction mixture before incubation. The reaction mixtures were resolved by electrophoresis through 6% polyacrylamide gel in 0.5× Tris-borate EDTA buffer at 100 V. The Cy5 signals were detected using WSE-6200H LuminoGraph II (ATTO).

### Transient Expression Assays with Arabidopsis Mesophyll Protoplasts

Transient transformation of Arabidopsis mesophyll protoplasts was carried out with slight adjustments from the protocol described (Yoo et al. 2007). To generate the *35S::NLS-GFP* construct, the coding sequence of GFP was amplified using PCR, incorporating primers that encompass the NLS sequence. The resulting fragment was subsequently cloned into the *HBT-HA-NOS* plasmid (Yoo et al. 2007). For the generation of the *35S::NLS-RVE8-GFP, RVE8* coding sequence was amplified using PCR, then inserted into the *35S::NLS-GFP* plasmid. For the −0.2 kb *pVIN3_U_I::LUC* construct, the sequence containing −0.2 kb *pVIN3*, 5′-untranslated region (UTR), the 1^st^ exon, 1^st^ intron, and the 2^nd^ exon of *VIN3* were amplified by PCR and cloned into the *LUC-NOS* plasmid (Hwang and Sheen 2001). Protoplasts were isolated from the leaves of SD-grown wild type plants (Col) using an enzyme solution comprising 150 mg Cellulase Onozuka™ R-10 (Yakult), 50 mg Maceroenzyme™ R-10 (Yakult), 20 mM KCl, 20 mM MES-KOH (pH 5.6), 0.4 M D-mannitol, 10 mM CaCl_2_, 5 mM β-mercaptoethanol, 0.1 g bovine serum albumin. For the protoplast transfection, reporter and each effector constructs were co-transfected into the protoplasts. 100 μg of each plasmid DNA was mixed with the isolated protoplasts in a transfection buffer containing 0.1 M D-mannitol, 50 mM CaCl_2_, and 20% (w/v) PEG. Luciferase activity in the protoplasts was quantified using the Luciferase Assay System (Promega) and a MicroLumat Plus LB96V microplate luminometer (Berthold Technologies).

### Statistical analysis

Statistical analyses were conducted using Microsoft Excel or GraphPad. For unpaired Student’s t-tests as well as one- or two-way analysis of variance (ANOVA), post hoc multiple comparison tests were employed. Significance levels are as specified in the legends of each figure, with p < 0.05 being deemed statistically significant.

## Resource availability

The accession number for the raw RNA-seq data obtained in this study is NCBI’s Sequence Read Archive (SRA) BioProject: PRJNA1087161.

## Acknowledgments

We would like to thank Dr. David Jackson (Cold Spring Harbor Laboratory, NY, USA) for kindly providing *cct8-1; gl1; pRbcS::GFP∼GL1∼KN1^C^* seeds and *pCCT8::CCT8-GFP* constructs and thank Dr. Stacy Harmer (University of California, Davis, CA, USA) for supplying the *rve3*, *rve4*, *rve5*, *rve6*, *rve8*, *rve468*, *rve34568* seeds. This work was supported by the Cooperative Research Program for Agriculture Science and Technology Development (No. PJ01315201 and PJ01315401), Rural Development Administration, Republic of Korea. This work was also supported by National Research Foundation of Korea (NRF) grants funded by the Korean government (MSIT) (No. 2019R1A2C2004313, 2021R1A5A1032428, and 2022R1A2C1091491). M. Jeon and D. Jeong were supported by Brain Korea 21 Plus Project. G. Jeong was also supported by the Stadelmann-Lee Scholarship Fund, Seoul National University, Seoul, Korea.

## Author Contributions

G.J., I.L. designed the research. G.J., M.J., J.K., D.J., J.S. and Y.S. performed the experiments. G.J, M.J, Y.S, D.H, Y.H. and I.L. analyzed the data. G.J. and I.L. wrote the paper. All authors read and approved the final manuscript.

